# Contrasting roles of Beclin-1 in pain hypersensitivity and anxiety-like behaviours in a mouse model of neuropathic pain

**DOI:** 10.1101/2023.12.18.572240

**Authors:** Fariya Zaheer, Gabriel J. Levine, Ana Leticia Simal, Paige O. Reid, Reza Fatemi, Tami A. Martino, Giannina Descalzi

**Affiliations:** Department of Biomedical Sciences; Center for Cardiovascular Investigations, Ontario Veterinary College, University of Guelph

## Abstract

Chronic pain is a debilitative disease affecting 1 in 5 adults globally^1^. The current understanding of chronic pain remains inadequate, coupled with few available therapeutics for the treatment of associated mental health disorders. Cellular homeostasis is crucial for normal bodily functions and investigation at the cellular levels may reveal a better understanding of the processes that occur leading to the development of chronic pain. Using the spared nerve injury (SNI) model of neuropathic pain, we found that adult male mice with impaired BECLIN-1 function show enhanced mechanical and thermal hypersensitivity compared to wildtype controls. Remarkably, we found that while SNI induced increases in anxiety-like behaviours in wildtype mice, this was not observed in mice with impaired BECLIN-1 protein function. Our data thus indicates that BECLIN-1 is differentially involved in the nociceptive and emotion related effects of chronic neuropathic pain.

## INTRODUCTION

Chronic neuropathic pain is a debilitating condition that is resistant to currently available treatments. Neuropathic pain develops upon direct and/or indirect injury to peripheral nervous tissue^2^. Mounting evidence has been reported indicating that long term potentiation (LTP) might be a critical mechanism promoting the development of chronic pain^3,4^. Several studies have also found the upregulation of N-methyl-D-aspartate (NMDA) receptors in the rodent models of neuropathic chronic pain, highlighting LTP as the source behind the hypersensitivity component of neuropathic pain^5–8^. Additionally, upon neuropathic injury, a change in cellular homeostasis is evident via the induction of neuroinflammation, which serves a protective role through the recruitment of immune cells to the site of injury to initiate damage control^9^. A key regulator of homeostasis is the process of autophagy which ensures appropriate disposal of damaged intracellular organelles to maintain equilibrium. BECLIN-1, a modulator of the PI3-K pathway, plays a major role in the process of macro-autophagy and is responsible for the delivery of autophagosomes to lysosomes for intracellular degradation, promoting autophagy and restoring cellular homeostasis^10,11^. Additionally, the BECLIN-1 protein drives autophagy to control the immune response initiated by microglia. Impaired function of the BECLIN-1 protein leads to continued immune response activation, causing excessive neuroinflammation through constant secretion of cytokines from microglia via the NLRP3 pathway, ultimately resulting in the development of neurodegenerative diseases^12^. Uncontrolled neuroinflammation could exacerbate the immune response contributing to the development of neuropathic pain. Notably, inflammation through the actions of prostaglandins and cytokines have been shown to impact molecular processes involved in LTP^13^. In this study, we used the spared nerve injury model of neuropathic pain to determine the role of BECLIN-1 in chronic pain development^14,15^. For our transgenic cell line, we used Bcl2^AAA^ mice^16^. These mice contain a knock in mutation in the B-cell lymphoma-2 (Bcl-2) protein, substituting hydrophobic alanine amino acids for phosphorylated amino acids at position T69A, S70A, S84A. The new alanine amino acids are unable to be phosphorylated so Bcl-2 stays bound to BECLIN-1, and the interaction cannot be dissociated, rendering the BECLIN-1 protein inhibited. Inhibition of BECLIN-1 protein prohibits it from performing a key role in autophagy^16^. We found that the Bcl2^AAA^ mice, which have an abnormal function of BECLIN-1 protein^16^, show enhanced SNI induced pain hypersensitivity when compared to wildtype mice.

Chronic pain is also often accompanied with mental health related discomfort, as it has been noted that over half of all people suffering from chronic pain also suffer from comorbid anxiety and/or depression^17^. One mechanism through which chronic pain may alter anxiodepressive states could be the enhanced inflammation levels of those with neuropathic pain^18^. Previous studies have linked inflammation with the development of anxiety and other emotional disorders. We found that Bcl2^AAA^ mice exposed to SNI behaved similarly in the elevated plus maze and open field tests as compared to sham treated controls. In contrast, wildtype mice exposed to SNI showed increased anxiety-like behaviours compared to their sham treated counterpart. Our data thus indicates that BECLIN-1 may prevent pain hypersensitivity, whilst promoting pain-induced increases in anxiety-like behaviours.

## METHODS

### Animals

We used male adult (11-12 week old) C57BL/6NCrl (Charles River) and male adult (11-12 week old) Bcl2^AAA^ knock-in mutation mice (Jackson Lab 018430) for all experiments. All mice were housed in threes in an animal facility with a 12-hour light-dark cycle (7:00 on, 19:00 off), in a room with an ambient temperature of 21-24° C with access to food and water provided *ad libitum.* All cages were provided with environmental enrichment, consisting of Crink-l’Nest bedding (Lab Supply, TX, USA), a Kimwipe, a polycarbonate igloo, and an elevated polycarbonate loft suspended from the cage roof. All procedures were approved by the Animal Care Committee at the University of Guelph.

### Anxiety-like behavioural tests

Elevated plus maze (EPM) and the open field test (OFT) were used to assess behaviour in all mice^19,20^. The elevated plus maze consists of two open arms (29cm), and two arms enclosed by walls (29cm). Mice were placed in the center of the EPM and allowed to freely explore for 5 minutes. Tests were recorded and analyzed using Noldus EthoVision XT software. Percent time spent exploring open arms (relative to total exploration of open and closed arms) were compared. Similarly, the open field tests consist of a white square box (45cmx45cm) in which mice are placed from one of the four corners and allowed to explore the field freely for 5 minutes. Percent time spent in the center (relative to total exploration of center and borders) were compared between cohorts. Experiments were performed by a researcher blind to treatment and genotype.

### Mechanical sensitivity test

Mechanical thresholds were determined through Von Frey filaments using the up-and-down method^21,22^. All mice were habituated to the testing apparatus which consisted of a plexiglass container, with an elevated wire mesh flooring for 2 hours for 2 days and baseline testing occurred on day 3. Following the surgery, Von Frey tests took place on day 3, 7, 14, 21 and 28 after habituating for 40 minutes on each testing day. Mechanical nociceptive thresholds were recorded after nociceptive responses were observed 50% of the time per stimulus. Experiments were performed by a researcher blind to treatment and genotype.

### Thermal sensitivity test

Thermal sensitivity to pain was assessed using the Hot Plate Tests (Bioseb, FL, USA) as previously reported^22^. Mice were placed on a plate heated to 50 degrees Celsius and the latency to withdraw, flick, or lick the hind paws was recorded. Each trial was limited to 30 seconds with 3 trials in total, divided with 10 minutes of rest period. The hot plate test took place pre- and post-op (day 29 after SHAM/SNI surgery). Experiments were performed by a researcher blind to treatment and genotype.

### Neuropathic Pain Model

The spared nerve injury (SNI) model was used as previously reported^14,15,20^. Briefly, mice were anesthetized using 1-3% isoflurane for surgery. Briefly, skin and muscle incisions were made on the left hindleg at midthigh level, revealing the sciatic nerve and its three branches. The common peroneal and tibial nerves were carefully ligated with 8.0 Vicryl suture (Ethicon, Johnson & Johnson Intl.), transected, and a 1- to 2-mm sections of each of these nerves were removed. The tibial nerve was left intact. Skin was then closed and stitched with 5.0 Prolene sutures (Ethicon, Johnson & Johnson Intl). Sham-operated mice were exposed to the same procedure, but all nerves were left intact and untouched.

### Statistics

All statistical analyses were performed using Graph Pad PRISM (ver. 10.1.2). Repeated measures (RM) two-way ANOVAs and unpaired student t-tests were used. Where appropriate, Holm-Sidak’s post-hoc analyses were employed.

## RESULTS

### Disruptions of Beclin-1 enhance mechanical allodynia caused by nerve injury

To investigate the role of the BECLIN-1 protein in relation to chronic neuropathic pain, we assessed the pain behaviour expressed by mice following the spared nerve injury. Prior to surgery, we investigated whether any baseline differences existed between the transgenic and wildtype mice. Following mouse handling for 5 days, baseline measures for mechanical and thermal pain sensitivity were conducted, and no differences were observed between transgenic and control cohorts (*Figure 1*). Next, mice were divided randomly into two groups to either be assigned for the neuropathic pain model (SNI) and for control (SHAM) within each cohort of Bcl2^AAA^ and wildtype mice. Following the SNI/SHAM surgery, we found that Bcl2^AAA^ mice exposed to SNI showed significant reductions in their mechanical thresholds and displayed significantly larger decreases in thresholds compared to male wildtype SNI mice (See *Table 1*). Both mouse lines subjected to surgical controls (SHAM) displayed no significant changes in mechanical thresholds (*Figure 1A*) when compared with their baseline. Additionally, as can be seen in *Figure 1B* and outlined in *Table 2*, the right hind paw of all SHAM and SNI mice remained unaffected following either SHAM or SNI procedures.

**Figure 1A,B.**
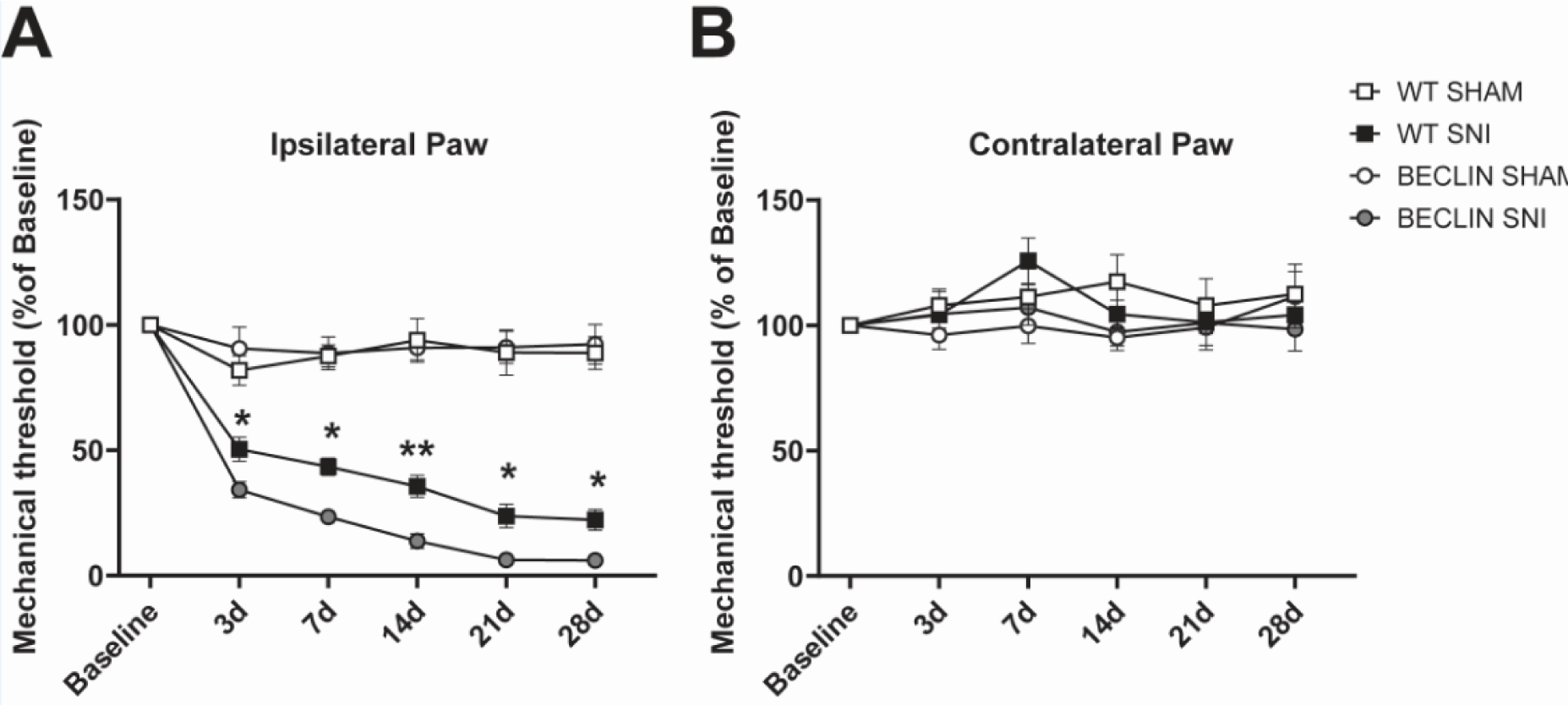
Chronic neuropathic pain causes mechanical allodynia in the SNI group for both wildtype and Bcl2^*AAA*^ transgenic cohorts. Bcl2^*AAA*^ knockouts following the spared nerve injury (SNI) in their left hind paw display increased hypersensitivity at a faster rate as compared to the wildtypes, significantly starting on day 3 up to until day 28. The right paw of all cohorts remained unaffected following both SNI and SHAM protocols post-treatment when compared to their initial baseline. Pain thresholds are recorded when pain responses are observed at 3 out of 5 applications of the Von Frey filament to the lateral aspect of the hind paw. Baseline tests were conducted before any treatment (SNI/SHAM). N=10-12/group, *P<0.05, **P<0.01.

**Table 1.** Von Frey thresholds for each cohort’s left paw (ipsilateral). Pain thresholds for left paw, conducted using the Von Frey test are outlined as mean, taken from percent of baseline for each mouse within a cohort, all averages were then divided to find group mean and standard error of the mean. All baselines are set as 100 from which each day of experiments are compared to.

| **Figure 1a**<br>**Left Paw** | **Mean thresholds in percent baseline (%) ± sem** |  |  |  |  |  | **Statistical analysis** |
| --- | --- | --- | --- | --- | --- | --- | --- |
| | Baseline | 3d | 7d | 14d | 21d | 28d | Two-way RM ANOVA<br>Time:<br>$F_{5,220}=68.83$,<br>$p<0.001$<br>TimeXgenotype:<br>$F_{15,220}=15.53$,<br>$p<0.001$<br>Genotype:<br>$F_{3,44}=95.13$,<br>$p<0.001$ |
| WT Sham | 100± 0.0 | 81.8±5.8 | 87.6±4.4 | 93.8±8.7 | 89.0±9.0 | 88.8±6.5 | |
| WT SNI | 100± 0.0 | 42.7±4.8 | 41.3±3.5 | 29.7±4.3 | 22.7±4.6 | 18.7±4.1 | |
| Bcl2<sup>AAA</sup> Sham | 100± 0.0 | 90.6±8.5 | 88.7±6.5 | 90.9±5.0 | 91.1±6.4 | 92.3±7.9 | |
| Bcl2<sup>AAA</sup> SNI | 100± 0.0 | 34.3±3.2 | 23.4±2.6 | 13.8±2.9 | 6.3±1.0 | 6.1±1.1 | |

**Table 2.** Von Frey thresholds for each cohort’s right paw (contralateral). Pain thresholds for right paw, conducted using the Von Frey test are outlined as mean, taken from percent of baseline for each mouse within a cohort, all averages were then divided to find group mean and standard error of the mean. All baselines are set as 100 from which each day of experiments are compared to.

| **Figure 1b**<br>**Right Paw** | **Mean thresholds in percent baseline (%) ± sem** |  |  |  |  |  | **Statistical analysis** |
| --- | --- | --- | --- | --- | --- | --- | --- |
| | Baseline | 3d | 7d | 14d | 21d | 28d | Two-way RM ANOVA<br>Time:<br>$F_{4.3,187.7}=1.511$,<br>$p=0.1973$<br>TimeXgenotype:<br>$F_{15,220}=0.9594$,<br>$p=0.4994$<br>Genotype:<br>$F_{3,44}=0.7816$,<br>$p=0.5106$ |
| WT Sham | 100± 0.0 | 108.0±6.4 | 111.3±11.2 | 117.4±10.8 | 107.8±10.8 | 112.5±11.9 | |
| WT SNI | 100± 0.0 | 104.3±5.4 | 125.8±9.0 | 104.5±5.5 | 101.2±4.5 | 104.3±5.4 | |
| Bcl2<sup>AAA</sup> Sham | 100± 0.0 | 96.2±5.7 | 99.8±6.9 | 95.1±3.3 | 99.2±9.0 | 111.5±10.0 | |
| Bcl2<sup>AAA</sup> SNI | 100± 0.0 | 104.5±9.0 | 107.2±9.1 | 97.3±7.3 | 101.0±9.0 | 98.5±8.6 | |

We also assessed the effects of SNI on thermal thresholds in wildtype and Bcl2^AAA^ mice using the hot plate test. At 29 days post SNI or SHAM surgery, mice were re-tested on the Hot Plate (50°C). When compared to baseline, both transgenic and wildtype mice in the SNI cohort showed a significant reduction in their latency to withdraw their paw on the heated plate (*Figure 2A,B*). Once again, Bcl2^AAA^ mice showed significantly larger reductions in their responses when compared to wildtype male mice in the SNI cohort (WT SNI: 11.5 ± 1.0 seconds post-op as compared to WT SNI pre-op: 19.3±1.3seconds; Bcl2^AAA^SNI post-op: 7.7 ±1.0 seconds from their baseline of 16.1±1.1seconds). The SHAM controls for both transgenic and wildtype mice showed no difference in their response when compared to their baseline measures conducted before the surgery (WT SHAM post-op: 18.6±1.3seconds from their baseline of 20.4±1.7seconds; Bcl2^AAA^ SHAM post-op 16.8±1.0seconds compared to their baseline of 16.1±1.2seconds; *Figure 2A,B*). Remarkably, as shown in *Figure 2*, Bcl2^AAA^ mice who received the SNI surgery showed significantly shorter latencies to withdraw their paw compared to Bcl2^AAA^ SHAM, wildtype SNI and wildtype SHAM.

**Figure 2A,B.**
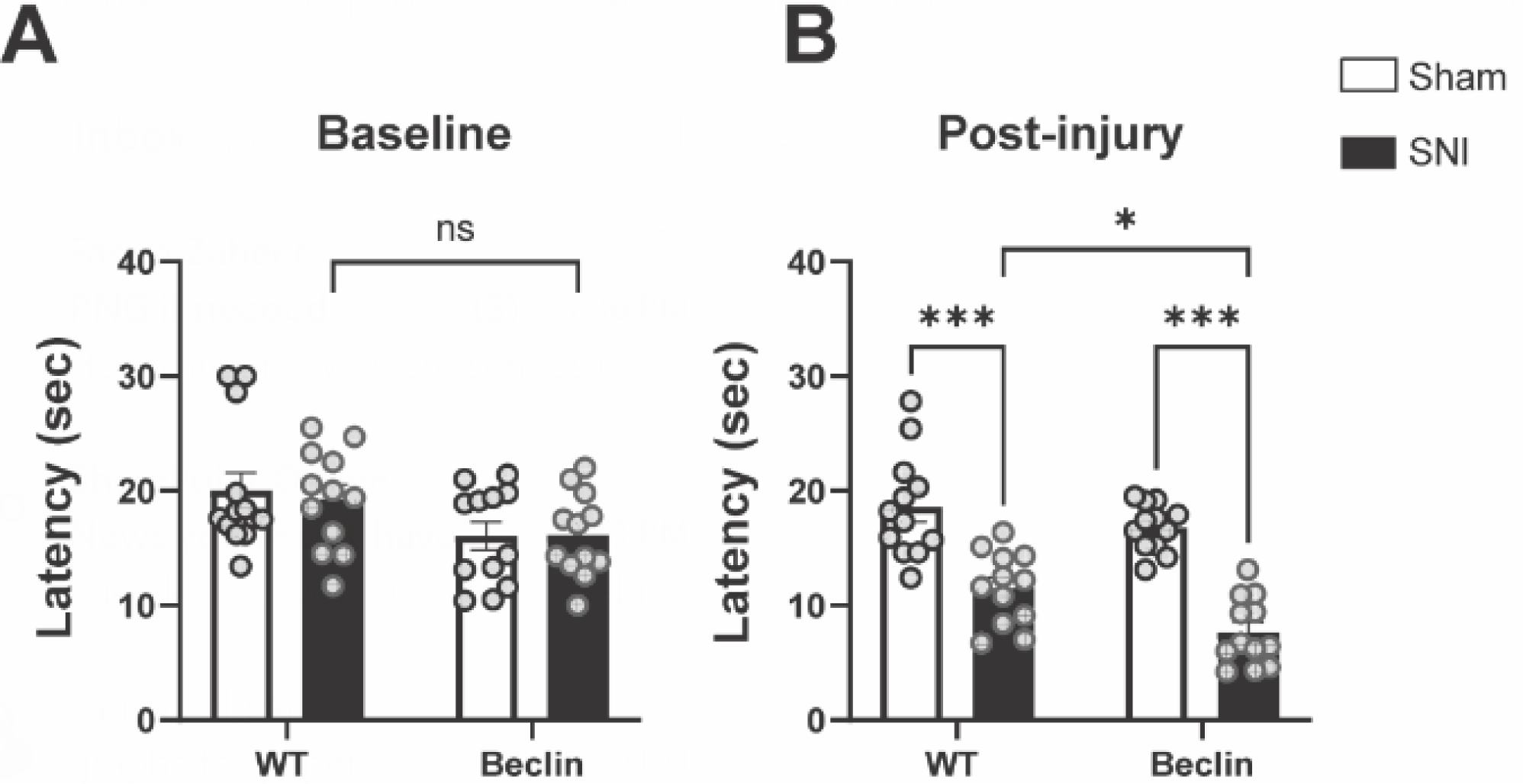
Both wildtype and transgenic Bcl2^*AAA*^ mice SNI cohorts display pain hypersensitivity when subjected to the hot plate test. SNI groups had a decreased latency to display nocifensive behaviour following surgery, while SHAM groups in both cohorts exhibited no significant changes when compared to their initial baseline values shown in (A). Significant differences were observed between SNI groups, as Bcl2^*AAA*^. Baseline tests were conducted before any SNI/SHAM treatment, post-injury measurements were collected on Day 29. n=12/group. *P<0.05, ***P<0.001.

### Disruptions of Beclin-1 does not affect SNI-induced changes in anxiety-like behaviours

Studies have linked inflammation with the unpleasant symptoms of anxiety and related mental health disorders in clinical settings^23–25^. We previously showed that chronic neuropathic pain increases anxiety-like behaviour in male mice^20,26^. We next sought to determine if disruptions in Beclin-1 signaling can affect SNI-induced increases in anxiety-like behaviours. To do this, we used the elevated plus maze and open field test on day 30-31, respectively, post-op. Surprisingly, on the elevated plus maze tests, we found that while WT SNI did exhibit increases in anxiety-like behaviours by spending less time in open arms, as shown in *Figure 3*, as compared to WT SHAM (WT Sham: 17.3± 2.8 %; WT SNI: 5.6 ± 1.3%). However, between the Bcl2^*AAA*^ cohorts, the mice assigned to SNI vs. SHAM cohorts did not show any preferences to either open or closed arms on the elevated plus maze (Bcl2^AAA^ Sham: 21.4 ± 2.6%; Bcl2^AAA^ SNI: 16.4±3.3%). Such results also translated to the open field test where exploratory behaviour is displayed through a rodent’s choice to spend time in the center of the open-field maze. Similar to the elevated plus maze test results, we found that the WT SNI cohort spent the shortest amount of time in the center as compared to their WT SHAM counterparts, indicating their anxious behaviour (*Figure 4*). As compared to WT, Bcl2^*AAA*^ treatment groups (SNI/SHAM) showed no preference for the center or the borders of the open field test (WT Sham: 22.3 ±3.7 %; WT SNI: 11.8±1.8%; Bcl2^AAA^ Sham: 17.5±1.9%; Bcl2^AAA^ SNI: 15.8±1.9%)

**Figure 3.**
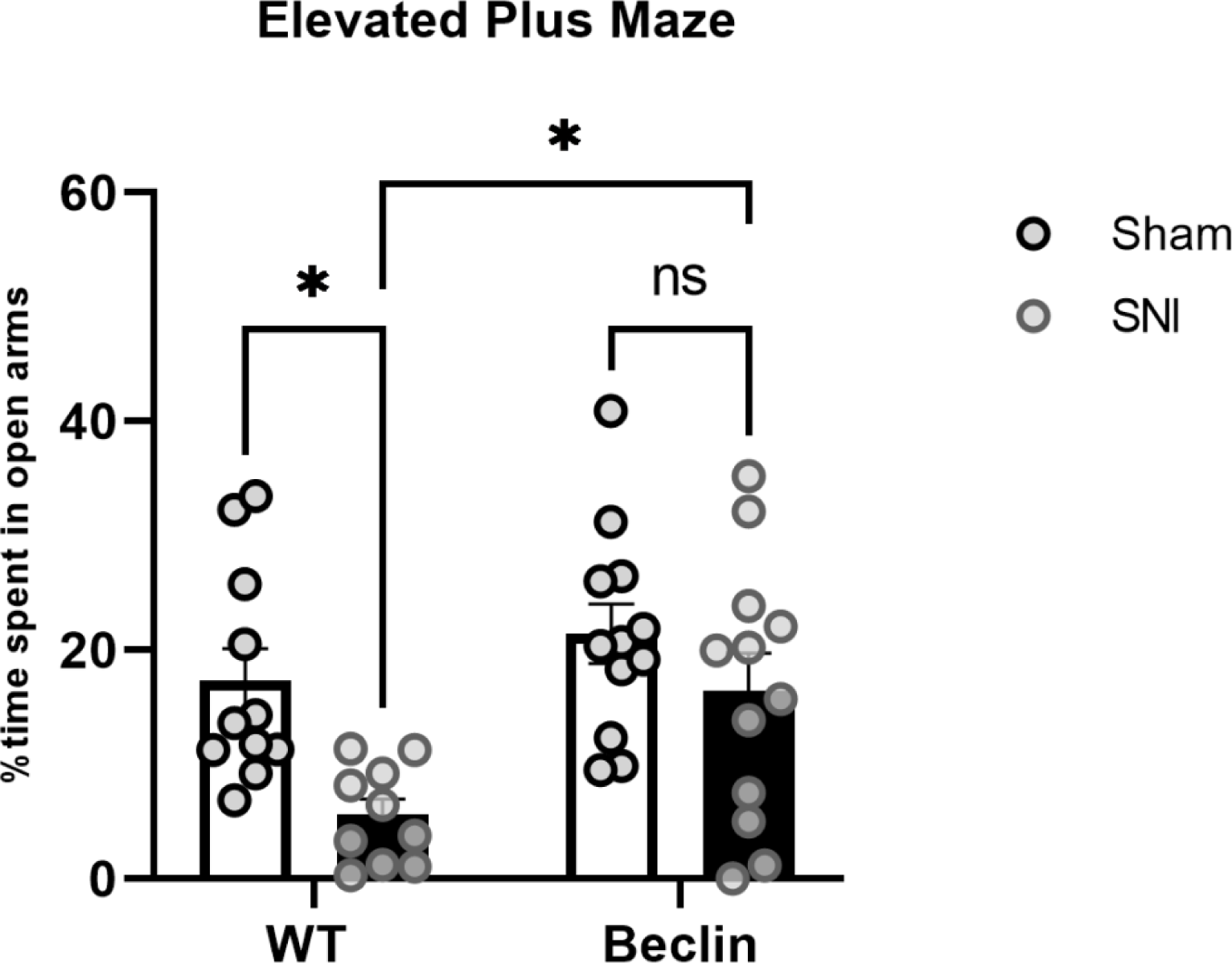
Wildtype SNI mice demonstrated more anxiety-like behaviour than Bcl2^*AAA*^ SNI mice, spending less time in the open arms when subjected to the elevated plus maze during a 5-minutes trial. The percent of time spent in the open arms was the percentage of the total time spent in the maze (closed+open arms). Time spent in an area is measured when all four paws of a mouse are contained within the zone. N=12/beclin SHAM and SNI. N=11 WT Sham. N=10 WT SNI, day 30-31 post-op SNI/SHAM. * P<0.05.

**Figure 4.**
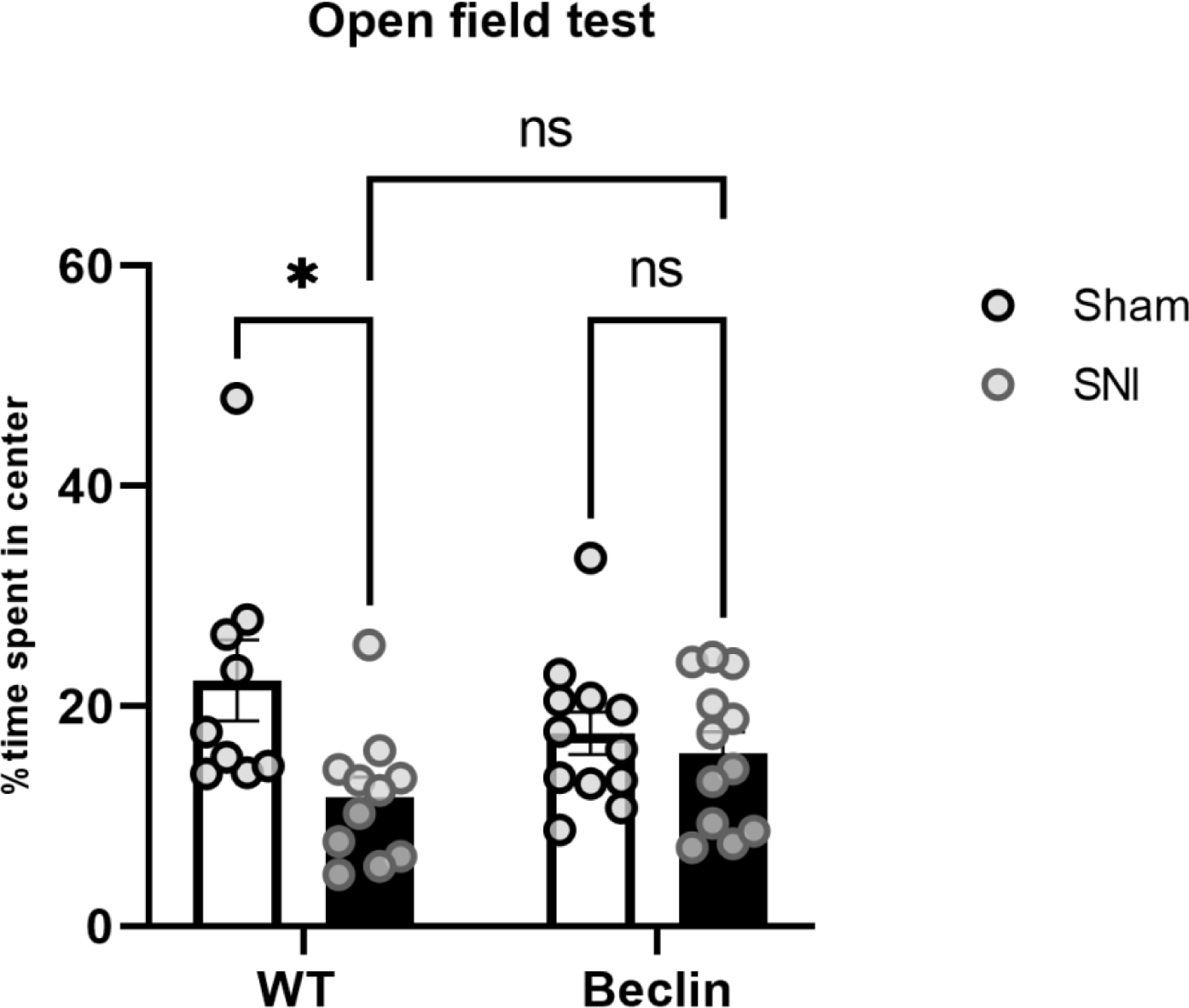
Wildtype SNI mice spend less time in the center of the open-field maze than wildtype SHAM mice, but both Bcl2^*AAA*^ mice cohorts (SNI/SHAM) showed no preference for the open vs. closed spaces. The percentage of time spent in the center of the five-minute trial was recorded. Time spent in the center was recorded when all four paws of the mice were within the central region of a 45 centimeter x 45 centimeter region of the open-field box. N=9/WT SHAM. N=11/WT SNI. N=12/ both Beclin SNI and SHAM, day 30-31 post-op SNI/SHAM. * p<0.05

We also confirmed that the neuropathic pain induced by using the spared nerve injury (SNI) or the control protocol (SHAM) did not impair a mouse’s motor ability and such observations were confirmed by collecting the distance on the elevated plus maze & open field tests (*Figure 5*;*Figure 6*).

**Figure 5.**
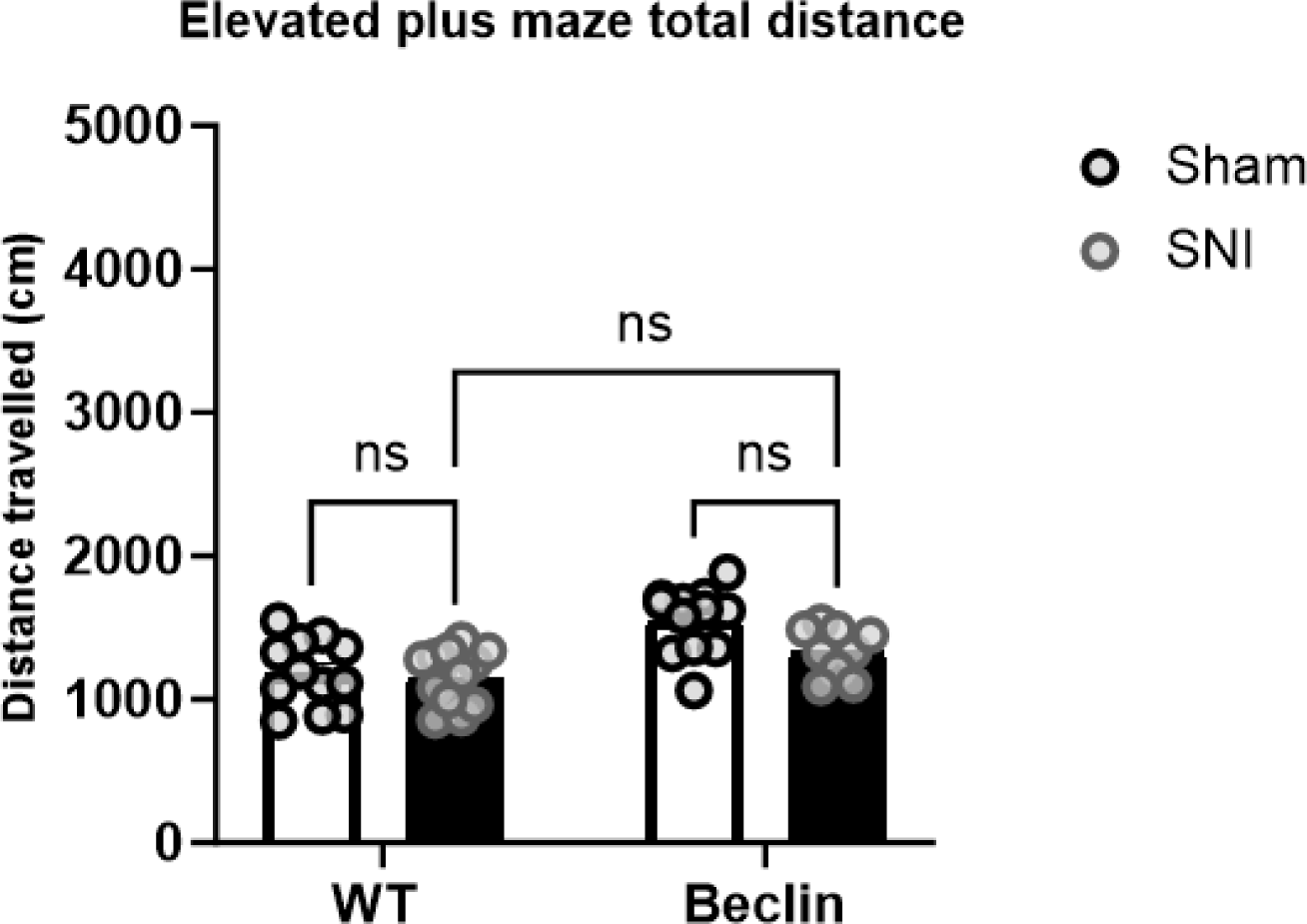
All groups travelled similar distances when subjected to the elevated plus maze test. Similar distance travelled during the elevated plus maze test for both Bcl2^*AAA*^ mice and wildtype suggests no differences in motor function between groups. N=10-12/group, day 30-31 post-op SNI/SHAM.

**Figure 6.**
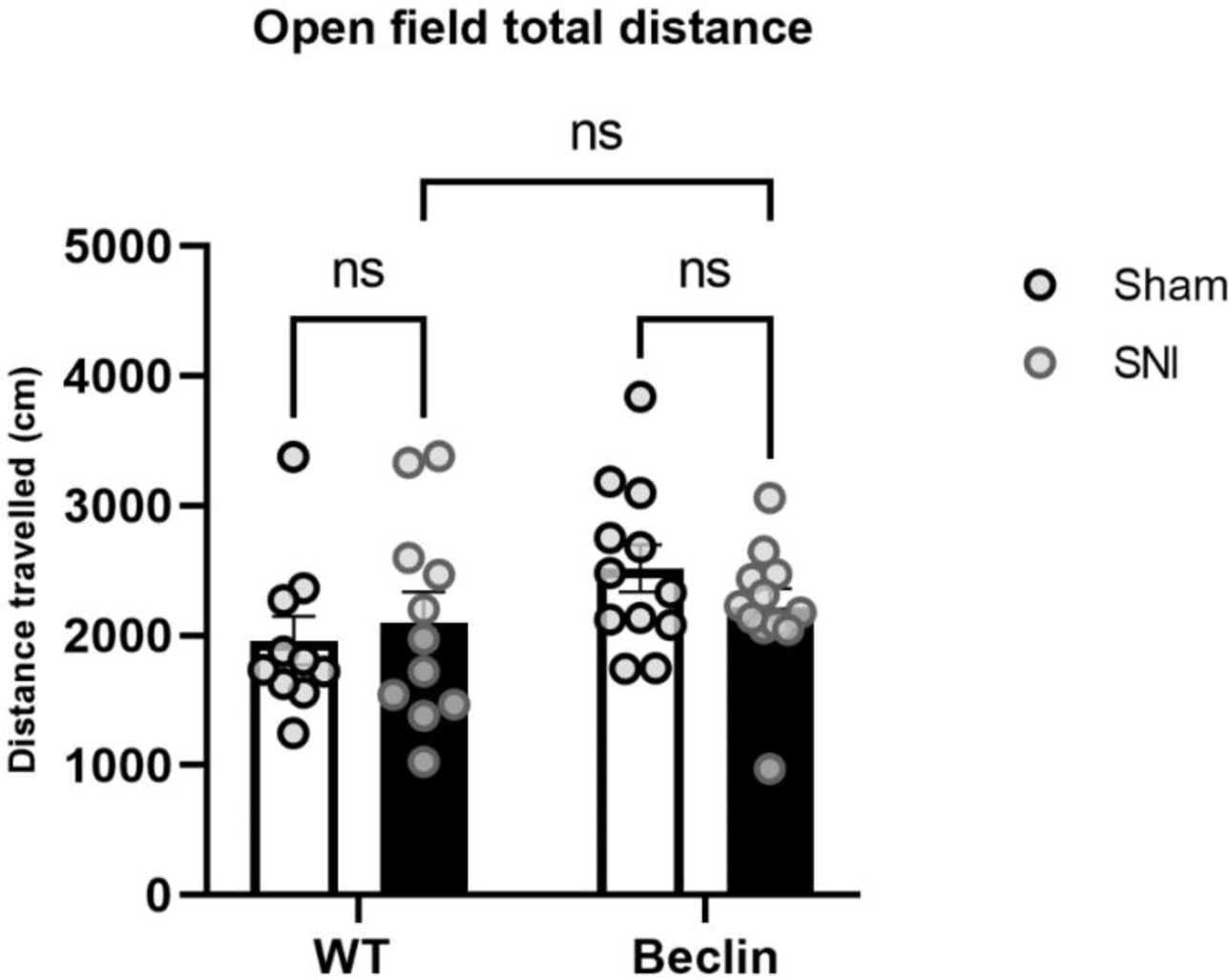
All cohorts demonstrated a similar distance travelled when subjected to the open-field test. Similar distance travelled in the open-field test for both Bcl2^*AAA*^ mice and wildtype suggests no differences in motor function between groups. N=10-12/group, day 30-31 post-op SNI/SHAM.

## DISCUSSION

In this study, we looked at whether the disturbance of stimulus induced autophagy would affect the development of chronic neuropathic pain by targeting the BECLIN-1 protein. The BECLIN-1 protein is an upstream protein of the PI3-K pathway and plays a crucial role in the formation of autophagosomes, which contain cargo destined to be degraded upon fusion with the lysosome. BECLIN-1 also plays a critical role in regulating the response of immune cells through inhibiting the production of certain pro-inflammatory cytokines at the gene expression level^27^. We believe that research towards developing a BECLIN-1 based intervention could achieve therapeutic relief for those struggling with chronic pain, as its reduced expression leads to enhanced pain hypersensitivity.

For our study, we began our experiments using male mice with a C57BL/6 background to test whether disturbing the autophagy pathway through inactivation of the BECLIN-1 protein leads to any pain related behavioural changes. Through the Von Frey filament test, we found that male mice with the Bcl2^AAA^ mutation who received SNI showed an enhanced pain response at a lower force compared to WT SNI cohorts significantly, starting from 3d post-op up until day 28. Bcl2^AAA^ mice exhibited pain responses from stimuli as low 0.07-0.04g, which translated to their pain response dropping from 100% down to 6% by day 28, confirming our hypothesis that mice with disturbances in the BECLIN-1 protein stimulus-induced autophagy pathway might be experiencing allodynia at a greater rate than the wildtype SNI cohort of male mice. While these mice are experiencing chronic neuropathic pain, their right paw showed no significant difference in their mechanical threshold upon application of Von Frey filaments, when compared to their initial baseline percentage. We believe that the enhanced response to low forces of filaments observed in Bcl2^AAA^ SNI mice is due to the decreased inflammatory response, as BECLIN-1 protein function is disrupted, impairing healing of the somatosensory injury induced by the SNI surgery.

We wanted to further investigate the pain response by testing for thermal hypersensitivity in each cohort. For this, we used the hot plate test, where a mouse is placed on a heated plate with a temperature of 50 degrees Celsius for a maximum of thirty seconds, and latency to exhibit a pain response is recorded. Pain responses on the hot plate test were recorded when mice raised or licked their paw. While both WT and Bcl2^*AAA*^ SNI cohort displayed hypersensitivity to pain by withdrawing their paw at a significantly faster rate than SHAM counterparts, Bcl2^*AAA*^ SNI cohort had an even greater reduction in their latency to withdraw their paws as compared to the WT SNI cohort. This further supports our conclusion that that while both WT and Bcl2^*AAA*^ SNI cohorts are experiencing neuropathic pain, it is the Bcl2^*AAA*^ cohort that is experiencing an enhanced nociceptive hypersensitivity.

Furthermore, we wanted to test if disruption of beclin-1 protein also affects pain-induced anxiety-like behaviours as chronic pain is often accompanied by mental health disorders in a clinical setting^17^. Inflammation in patients with chronic neuropathic pain has been linked to the onset of anxiety and other mental health-related symptoms^28^. We wanted to test if the transgenic mice line, who have been mutated to have reduced autophagy and possibly decreased inflammation, would show us such anxiety-like symptoms at a greater level than the wildtypes with SNI model of chronic neuropathic pain when subjected to experiments. We assessed such symptoms by using tests such as the elevated plus maze and the open field test which analyzes a mouse’s exploratory behaviour through their movement in certain (open/closed) zones, allowing us to equate the exploratory behaviour to determine their anxiety-like behaviours^19^. We found that while WT SNI did show enhanced anxiety-like behaviours compared to the WT SHAM cohort, the Bcl2^*AAA*^ cohort with abnormal BECLIN-1 protein function displayed no differences in anxiety-like behaviours. It is possible that impaired BECLIN-1 function results in a reduction of the inflammatory response due to decreased autophagy in comparison to wildtype^12,16,29^. If that is the case, our results may indicate that in response to peripheral nerve injury, such as caused by SNI, increases in inflammation promote a reduction of pain hypersensitivity; whereas they conversely cause increases in anxiety-like behaviours. Indeed, our findings show that Bcl2^*AAA*^ mice exposed to SNI did not develop increases in anxiety-like behaviour which are reliably seen in WT mice exposed to SNI^30,31^, whereas they did show enhanced pain hypersensitivity.

These findings could be attributed to the reduction in stimulus-induced autophagy and a diminished inflammatory response brought on by the immune system upon injury^32,33^. If the function of BECLIN-1 protein is interrupted, then this might serve as one of the many factors of enhanced pain sensitivity experienced by those struggling with chronic neuropathic pain. The role of BECLIN-1 protein extends beyond the pain hypersensitivity as it also seems to play a critical role in the affective component of pain. This suggest that a possible bidirectional role of the inflammatory response in chronic pain conditions, as while it may be protective against the enhanced nociceptive signalling from pain stimuli, it negatively affects the well-being of those with chronic neuropathic pain through its impact on the affective component of pain^34^. Such findings should be further explored through future investigation on the role of inflammation and pain hypersensitivity and whether enhanced or diminished inflammatory responses serves a protective role against the onset of mental health disorders brought on by the affective component of pain.

In conclusion, disruption of the BECLIN-1 protein increases nerve injury induced pain hypersensitivity, whilst conversely reducing pain-induced anxiety-like behaviour. This finding suggests that BECLIN-1 may have a protective role against pain hypersensitivity. This knowledge paves the way for the development of effective therapeutics that do not diminish the immune response while also preventing emotional discomfort.

